# S1-seq assay for mapping processed DNA ends

**DOI:** 10.1101/189662

**Authors:** Eleni P. Mimitou, Scott Keeney

## Abstract

During meiosis, the specialized cell division giving rise to gametes, numerous DNA doublestrand breaks (DSBs) are introduced at multiple places throughout the genome by the topoisomerase-like protein Spo11. Homologous recombination, a highly-conserved DSB repair pathway, is employed for their repair and ensures the formation of chiasmata and the proper segregation of homologous chromosomes. In the initial steps of recombination, end resection takes place, wherein Spo11 is endonucleolytically released and the 5′-terminal strands of each DSB are exonucleolytically processed, exposing the ssDNA necessary to identify a homologous repair template. DNA removed by DSB processing is reconstituted by DNA synthesis, which copies genetic information from the intact homologous template. We developed a next-generation sequencing assay, termed S1-seq, to study DNA end resection genome-wide at high spatial resolution during yeast meiotic recombination. The assay relies on the fact that removal of the ssDNA tails of resected DSBs marks the position where resection stopped. Molecular features of resection are revealed by sequencing of these ssDNA-to-dsDNA junctions and comparison to high-resolution Spo11 DSB maps. We describe the experimental and computational methods for S1-seq as applied to meiosis in the SK1 strain of budding yeast *Saccharomyces cerevisiae*, and discuss how it can also be applied to map DSBs and recombination intermediates.

## 1. Introduction

Meiotic homologous recombination plays a pivotal role in sexual reproduction and genetic diversity by promoting proper segregation of homologous chromosomes and by disrupting linkage groups (Hunter, 2015; Thacker, 2016). The event that initiates exchange of genetic information between parental chromosomes is the formation of DSBs by the transesterase Spo11 (Lam & Keeney, 2014)(Figure 1). These DSBs tend to cluster in narrow regions called hotspots (de Massy, 2013; Petes, 2001). Following break formation, the DNA ends undergo resection, the 5′→3′ degradation that gives rise to 3′ single stranded DNA (ssDNA) overhangs (Mimitou & Symington, 2009). This step is important to reveal the sequence proximal to the break, which when coated with the strand exchange proteins Rad51 and Dmc1 can invade the repair template in the homologous chromosome and initiate repair DNA synthesis, thus restoring the information lost during resection (Neale & Keeney, 2006). Shortly after the DSB is formed, the highly conserved Mre11-Rad50-Xrs2 (MRX) complex provides the Mre11 nuclease, which cooperates with Sae2 to nick the Spo11-bound DNA strands near the break (Cannavo & Cejka, 2014; Keeney, Giroux, & Kleckner, 1997; Moreau, Ferguson, & Symington, 1999; Neale, Pan, & Keeney, 2005). Mre11 then degrades the nicked strand towards the break via its 3′—5′ exonuclease activity, releasing Spo11 bound to short oligos (Garcia, Phelps, Gray, & Neale, 2011; Zakharyevich et al., 2010). The Mre11 nicks also constitute entry points for the 5′—3′ exonuclease, Exo1, which extends resection to an average of 822 nt (Mimitou, Yamada, & Keeney, 2017; Zakharyevich et al., 2010). During recombination in vegetative cells, a parallel pathway resects DSBs redundantly with Exo1, and depends on the concerted action of the Sgs1-Top3-Rmi1 complex and the Dna2 endonuclease (Mimitou & Symington, 2008; Zhu, Chung, Shim, Lee, & Ira, 2008).

**Figure 1.**
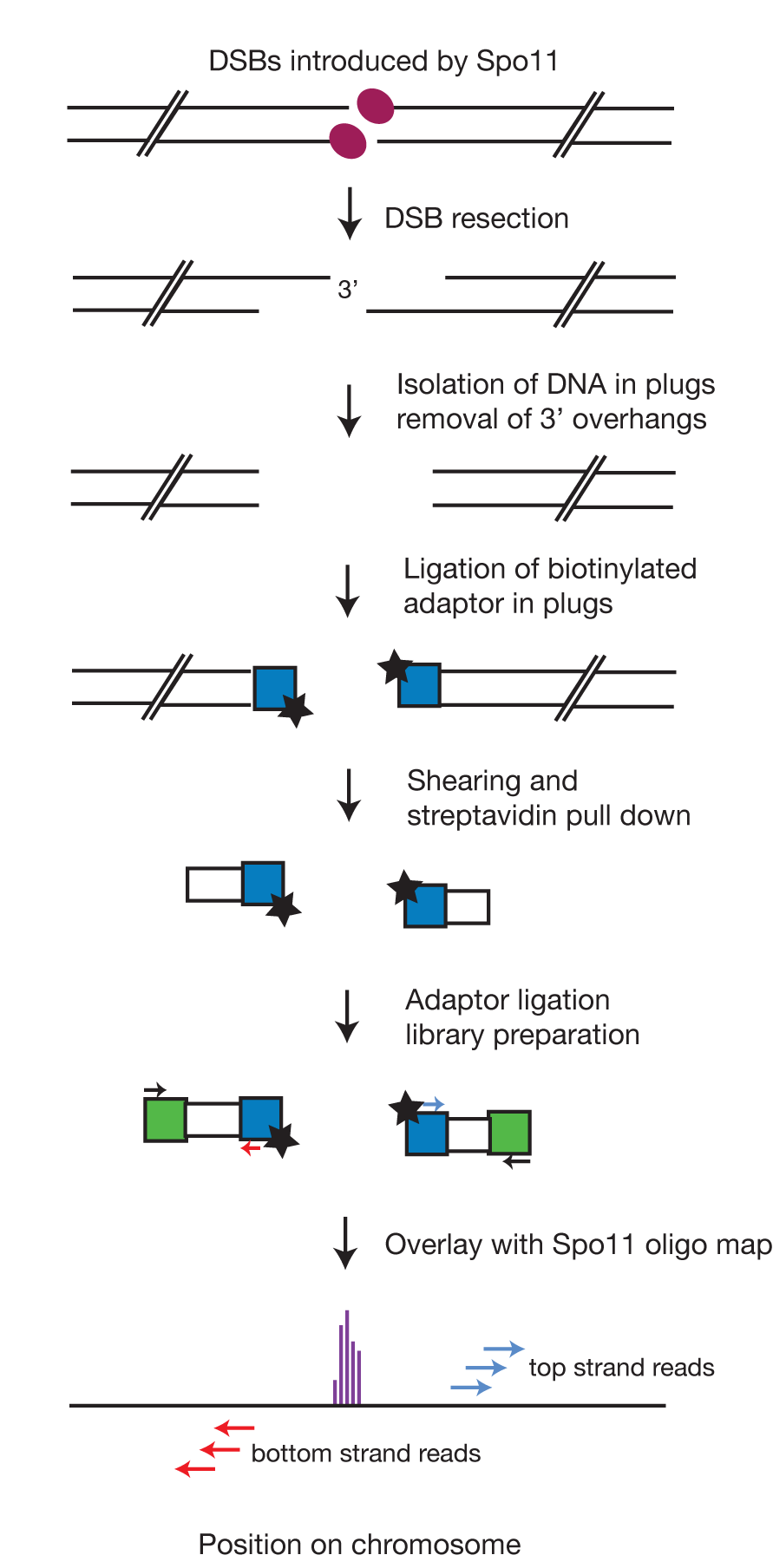
Schematic of the key steps employed in S1-seq: DNA from meiotic cells bearing processed Spo11 DSBs is embedded in agarose plugs to protect from shearing. Treatment with S1 nuclease and T4 DNA polymerase removes the 3′ overhang and prepares the ends for ligation with the 5′ biotinylated adaptor. Following extraction from the plugs and sonication, the biotinylated fragments are bound on streptavidin beads and the second adaptor is ligated. Low-cycle amplification by PCR creates the library that is submitted for next generation sequencing and eventually mapped to the reference genome.

## 2. Overview of S1-seq

The sensitivity of the 3′ ssDNA of resected DSBs to ssDNA-specific endonucleases provides a tool to map resection endpoints by sequencing the region where ssDNA transitions to dsDNA (Figure 1) (Mimitou et al., 2017). We isolate genomic DNA of meiotic cells in low-melting point agarose plugs to protect the DNA from random shearing. The plugs are treated with S1 nuclease and T4 DNA polymerase, then the blunt-ended chromosome fragments are ligated to a 5′ biotinylated dsDNA adaptor. Following extraction of DNA from the agarose plugs and sonication, we affinity purify the biotinylated fragments and ligate different adaptors to the opposite ends. The DNA is then amplified by low cycle PCR and deep sequenced, then the reads are mapped to a reference genome. Overlay of the sequencing reads with mapped Spo11 DSB sites (Pan et al., 2011) reveals features of resection genome-wide (Figure 1).

At least two biological replicates for each background and time point should be sequenced. In our hands, the assay displays high reproducibility (typical Pearson’s *r* = 0.98) (Figure 2A,B). Several features of the data confirm S1-seq as a readout of bona-fide resection endpoints. First, each DSB hotspot gives rise to surrounding reads that have the expected polarity: resection to the right results in endpoints that map to the top strand, whereas resection to the left gives rise to endpoints that map to the bottom strand (Figures 3A,B). Second, premeiotic samples or samples from the catalytically inactive *spo11-YF* mutant lack enrichment of signal around hotspots, confirming that the S1-seq signal is DSB-dependent (Figure 3B). Third, S1-seq read counts correlate well with Spo11-oligo counts (Figure 3C). Finally, the S1-seq signal displays the proper genetic control: it clusters more closely to hotspot midpoints in an *exol* nuclease-defective mutant and spreads further away in the *dmc1Δ* mutant, in which DSBs experience hyperresection (Bishop, Park, Xu, & Kleckner, 1992) (Figure 3D). S1-seq therefore is a selective, sensitive, and quantitative measure of DSB resection tract distributions. It should also be feasible to apply it to create high-resolution resection tract maps in other settings or systems.

**Figure 2.**
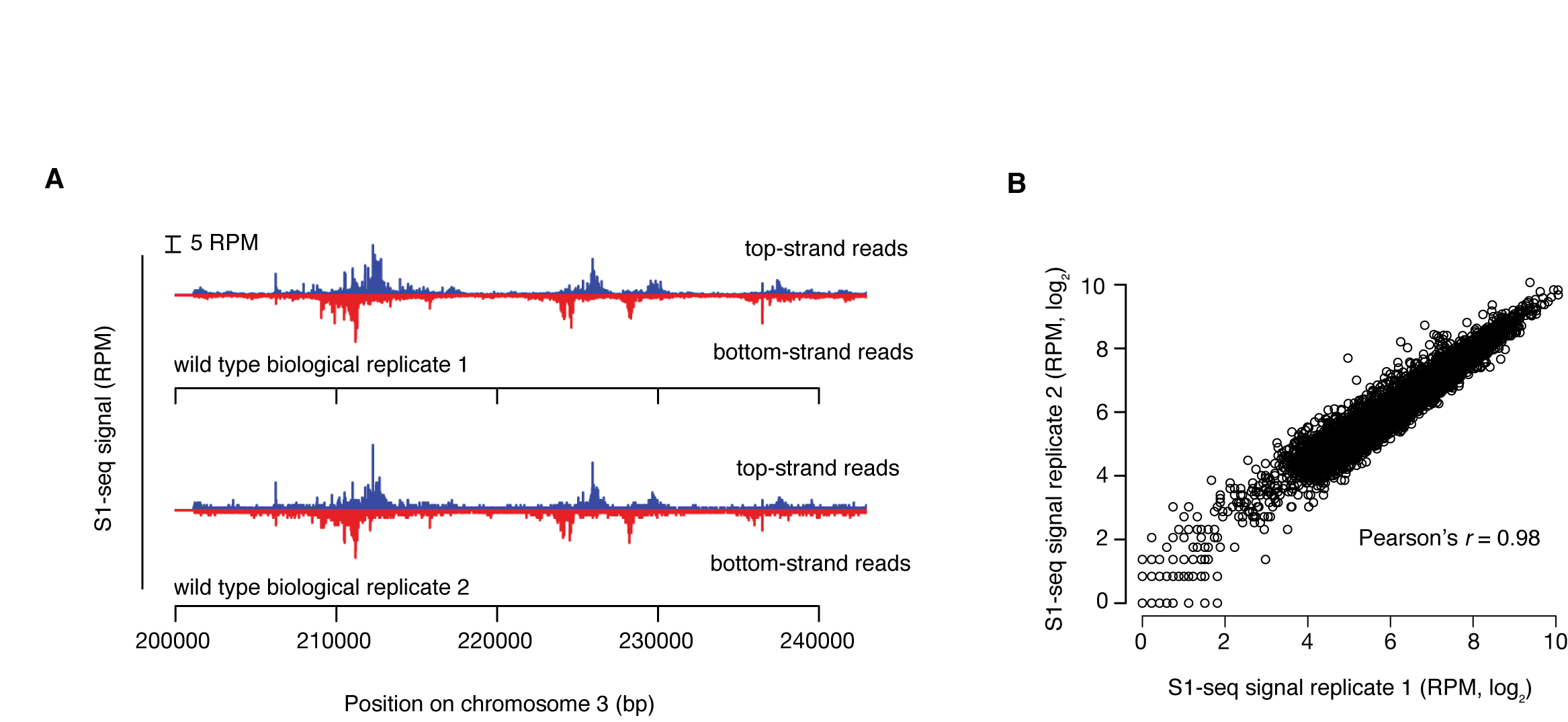
Reproducibility of S1-seq: A. Spatial reproducibility of S1-seq maps. Uniquely mapped S1-seq reads were normalized to reads per million (RPM). A region of chromosome III is shown as an example of the reproducibility between two wild-type datasets. B. Quantitative reproducibility of S1-seq maps. Normalized, uniquely mapped reads were summed in 1-kb nonoverlapping windows for two wild-type biological replicates. Adapted from Mimitou & Keeney, 2017.

**Figure 3.**
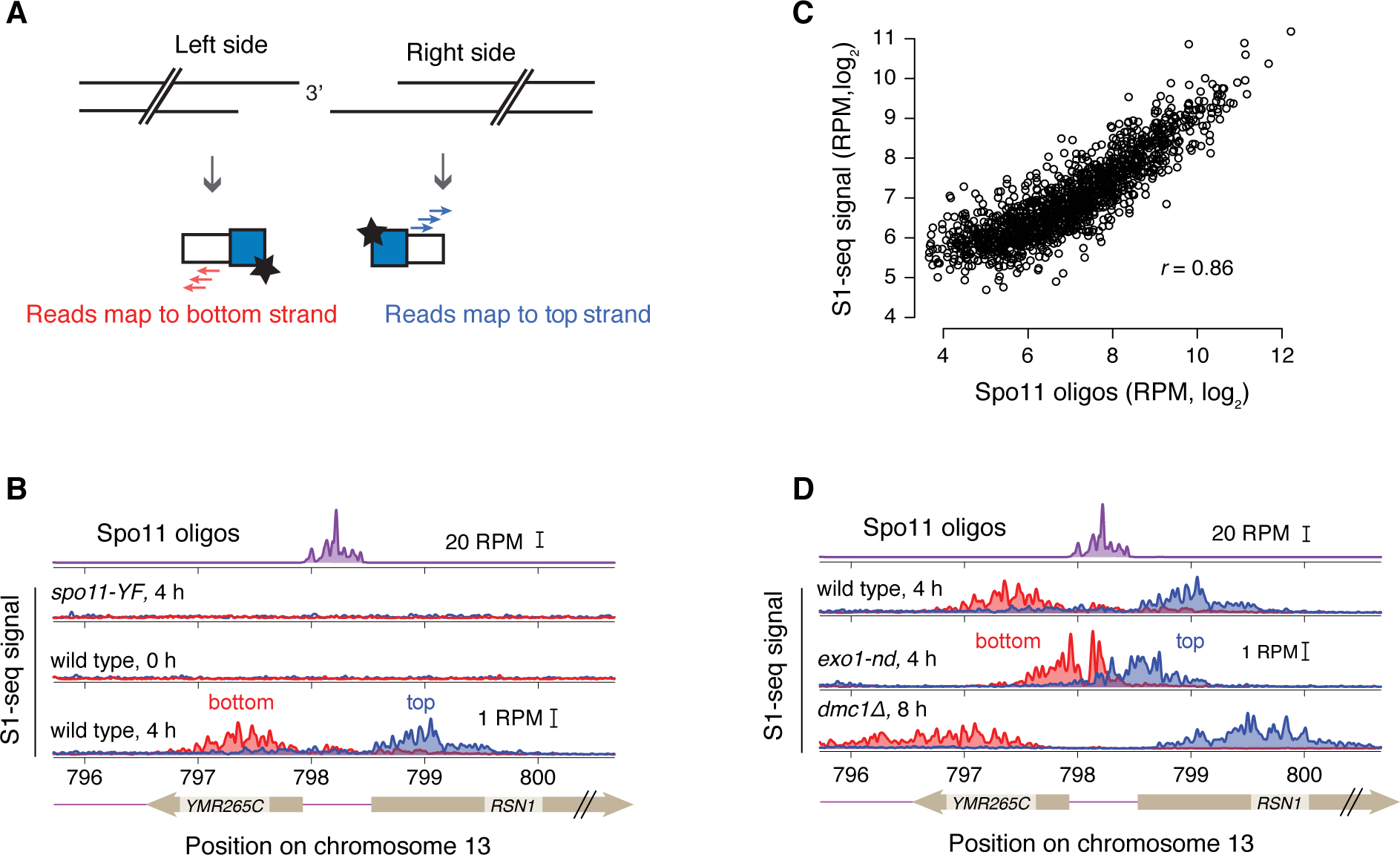
Validation of S1-seq: A. Schematic of the S1-seq signal polarity. Resection to the right or left should yield reads mapping to the top or bottom strands, respectively. B. An example of Spo11-oligo and strand-specific S1-seq reads (51-bp smoothing) surrounding the *RSN1* hotspot. C. S1-seq correlates well with Spo11 oligos: S1-seq signal (wild-type, 4 h) was summed from +200 to +1600 (top strand) and ‐200 to ‐1600 (bottom strand) relative to hotspot midpoints and plotted against the sum of the Spo11-oligo hitcount for each corresponding hotspot. For this analysis, hotspots with no other hotspot within 2 kb were used. D. Spo11-oligo and strand-specific S1-seq reads as in panel B. Adapted from Mimitou & Keeney, 2017.

## 3. Adaptations of S1-seq

### 3.1 Mapping unresected DSBs

We have shown that the sequential treatment with S1 nuclease and T4 DNA polymerase can degrade the 2-nt 5′ overhang of unresected DSBs to which Spo11 is still covalently bound (Mimitou et al., 2017). We therefore reasoned that S1-seq can provide a facile method for nucleotide-resolution mapping of the unresected Spo11 DSBs that accumulate in *sae2Δ* or *rad50S* mutants (Alani, Padmore, & Kleckner, 1990; McKee & Kleckner, 1997; Prinz, Amon, & Klein, 1997). As a proof of principle, we performed S1-seq in *sae2Δ* mutants, and observed DSB patterns similar to those revealed by traditional Spo11-oligo mapping (Figure 4A). For DSB mapping, S1-seq can be applied without any modification to the protocol below, as long as it is performed using cells of *sae2Δ* or *rad50S* background.

**Figure 4.**
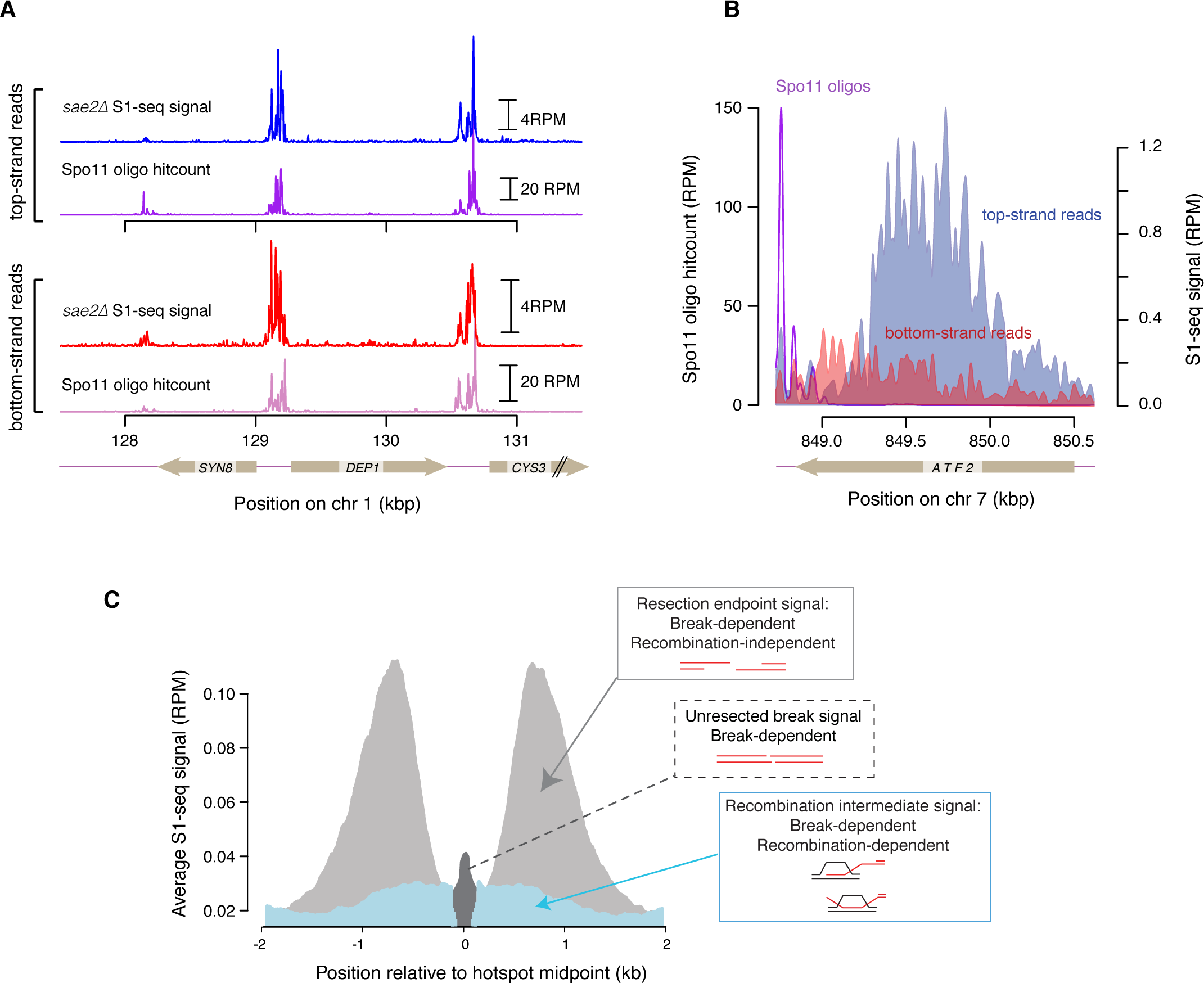
Adaptations of S1-seq: A. S1-seq in a *sae2Δ* background can reveal unresected Spo11 DSBs that correlate well with DSBs mapped by traditional Spo11-oligo sequencing. B. A snapshot of top-and bottom-strand S1-seq reads on the right side of a Spo11 hotspot. The most abundant top-strand reads represent resection endpoints whereas the bottom-strand signal originates from RIs. C. A graphical representation of the different species detected by S1-seq around hotspots: resection endpoints (light gray), RIs (blue), and unresected DSBs (dark gray). Adapted from Mimitou & Keeney, 2017.

### 3.2 Mapping recombination intermediate branchpoints

Along with resection signal, there is a weaker S1-seq signal with the “wrong” polarity, e.g., top-strand reads mapping to the left of hotspots and bottom-strand reads mapping to the right of hotspots (Figure 4B). As shown previously (Mimitou et al., 2017) this signal is Dmc1 dependent and derives from S1-sensitive recombination intermediates (RIs), probably displacement (D) loops from strand exchange. It is therefore possible to implement S1-seq toward the study of the spatial, temporal, and quantitative features of RI branchpoints by filtering S1-seq maps for reads of polarity opposite to resection-derived reads. A schematic summarizing the species detected by S1-seq is presented in Figure 4C.

## 4. Genomic DNA isolation

### 4.1 Yeast culture

Cells of desired genotype are induced into the meiotic program by nitrogen starvation and samples are withdrawn to follow DSB formation and processing.

#### 4.1.1 Material, Solutions, and Reagents

1. YPD plates: 1% Bacto yeast extract, 2% Bacto peptone, 2% dextrose, 2% Bacto agar. Autoclave.
2. YPD: 1% Bacto yeast extract, 2% Bacto peptone, 2% dextrose. Autoclave.
3. YPA: 1% Bacto yeast extract, 2% Bacto peptone, 2% potassium acetate, 0.001% antifoam. Autoclave.
4. SPM: 2% potassium acetate, 0.001% antifoam. Autoclave.
5. 2.8 L wide-mouth baffled Fernbach flasks.
6. 50 mM EDTA pH 8.0.
7. 15 mL conical tubes

#### 4.1.2 Procedure

1. Streak diploid cells from a frozen stock onto YPD plates and allow to grow at 30°C until single colonies are visible (approximately 36–48 h).
2. Inoculate a single colony into 20 mL liquid YPD medium and grow for >24 h at 30°C.
3. Using a spectrophotometer, measure cell density (OD_600_) of a 1:10 dilution of the overnight culture. Subculture into the appropriate volume of YPA medium to OD_600_ of 0.2 and grow in 2.8 L baffled flasks at 250 rpm at 30°C for 14 h.
4. Harvest cells by centrifuging at 2000 *g* for 3 min at 4°C and wash once with sporulation medium.
5. Resuspend cells in SPM in a 2.8 L baffled flask at an OD_600_ of 6.0 and incubate at 30°C, 250 rpm. Harvest cells at indicated time points: 75 mL for 0 and 2 h, 66 mL for 4 h, 60 mL for 6 h and 54 mL for 8 h. See note 1: Remove small volume samples to monitor progression of cell division by DAPI staining.
6. Wash cells with one half volume of 50 mM EDTA pH 8.0 and harvest by centrifugation.
7. Wash cells with 10 mL 50 mM EDTA and move into 15 mL falcon tubes
8. Harvest cells by centrifugation, remove the liquid, freeze on dry ice, and store at ‐80°C until plug preparation.

### 4.2 Isolation of DNA in plugs

Isolation of genomic DNA from the yeast cell samples is performed while the cells are embedded in low melting point (LMP) agarose to yield intact high molecular weight DNA.

#### 4.2.1 Material, Solutions, and Reagents

1. 500 mM EDTA pH 8
2. 10 mM Tris-HCl pH 7.5
3. 2-Mercaptoethanol
4. SCE solution: 1 M sorbitol, 0.1 M sodium citrate, 0.06 M EDTA, pH 7
5. Zymolyase 100T (US Biological)
6. LMP agarose (Lonza SeaPlaque)
7. Agarose plug molds (Biorad)
8. RNAse A 10 mg/mL (Thermo Scientific)
9. 20% SDS
10. Proteinase K, recombinant, PCR grade (Roche)
11. Glycerol
12. Solution 1: 5% 2-Mercaptoethanol, 1 mg/mL Zymolyase 100T in SCE solution
13. Solution 2: 0.45 M EDTA pH 8, 0.01 M Tris-HCl pH 7.5, 7.5% 2-Mercaptoethanol, 10 μg/mL RNase A
14. Solution 3: 0.25 M EDTA pH 8, 0.01 M Tris-HCl pH 7.5, 1% SDS, 1 mg/mL Proteinase K
15. Plug storage solution: 0.05 M EDTA pH 8, 50% glycerol
16. Low-retention filter tips
17. 15 mL conical tubes

#### 4.2.2 Procedure

1. Prepare fresh 2% LMP in 125 mM EDTA pH 8 and keep at 65°C.
2. Prepare fresh Solution 1 and keep at 40°C.
3. Resuspend each pellet in 150 μL 50 mM EDTA pH 8 and allow to thaw at 40°C.
4. Add 102 μL Solution 1 to the cells and mix well, so no clumps are present. See note 2.
5. Add 498 μL 2% LMP agarose, mix well, and quickly pipet into plug molds (~90 μL each).
6. Allow to solidify at 4°C for ≥30 min.
7. Express agarose plugs (3 at a time) into 3 mL of Solution 2 in 15 mL conical tubes and incubate at 37°C for 1 h.
8. Carefully aspirate Solution 2 and replace with 3 mL of Solution 3.
9. Incubate at 50°C overnight.
10. Wash plugs with 3 mL 50 mM EDTA pH 8, three times for 20 min each, with gentle shaking.
11. Aspirate last wash and add 3 mL plug storage solution.
12. Store at ‐20°C until all plugs are ready to be processed for library preparation.

## 5. Library preparation

### 5.1 Plug processing for overhang removal and adaptor ligation

All DNA treatments in this step are performed in the plugs to avoid pippeting and to minimize non-specific shearing of genomic DNA. For efficient in-plug overhang removal and adaptor ligation, a sequential treatment with S1 nuclease and T4 DNA polymerase is required before ligation. Even though S1 nuclease treatment is sufficient to remove the ssDNA tails, the subsequent ligation step is not efficient unless a clean-up reaction with T4 DNA polymerase is performed (E.P.M., unpublished observations).

Optional step: Some processed plugs can be set aside to serve as a control reaction, wherein a non-biotinylated P5 adaptor is ligated to the blunt ends after S1/T4 treatment. This reaction is then processed throughout the protocol in parallel with the biotinylated samples and serves to monitor background from non-specific binding of DNA to the beads.

#### 5.1.1 Material, Solutions, and Reagents

1. 1× TE: 10 mM Tris, 1 mM EDTA
2. 500 mM EDTA pH 8
3. 10× S1 buffer: 500 mM sodium acetate (pH=4.5), 2.8 M NaCl, 45 mM ZnSO_4_
4. S1 Nuclease (Promega)
5. 10 mg/mL (100×) BSA (NEB, currently discontinued, can be replaced with 20 mg/mL BSA)
6. 100 mM dNTPs set (Sigma Aldrich)
7. T4 DNA polymerase reaction buffer: 1 × T4 DNA ligase buffer, 1 × BSA, 100 μM dNTPs (each)
8. T4 DNA polymerase (NEB)
9. 10× T4 ligase reaction buffer (NEB)
10. T4 DNA ligase, 2000 U/μL (NEB)
11. P5-top oligo 100 μM, dissolved in 10 mM Tris-HCl pH 7.5, 50 mM NaCl, 1 mM EDTA pH 8 (see table 1)
12. P5-bottom oligo 100 μM, dissolved in 10 mM Tris-HCl pH 7.5, 50 mM NaCl, 1 mM EDTA pH 8 (see table 1)
13. Low-retention filter tips
14. 2 mL conical tubes

**Table 1:**
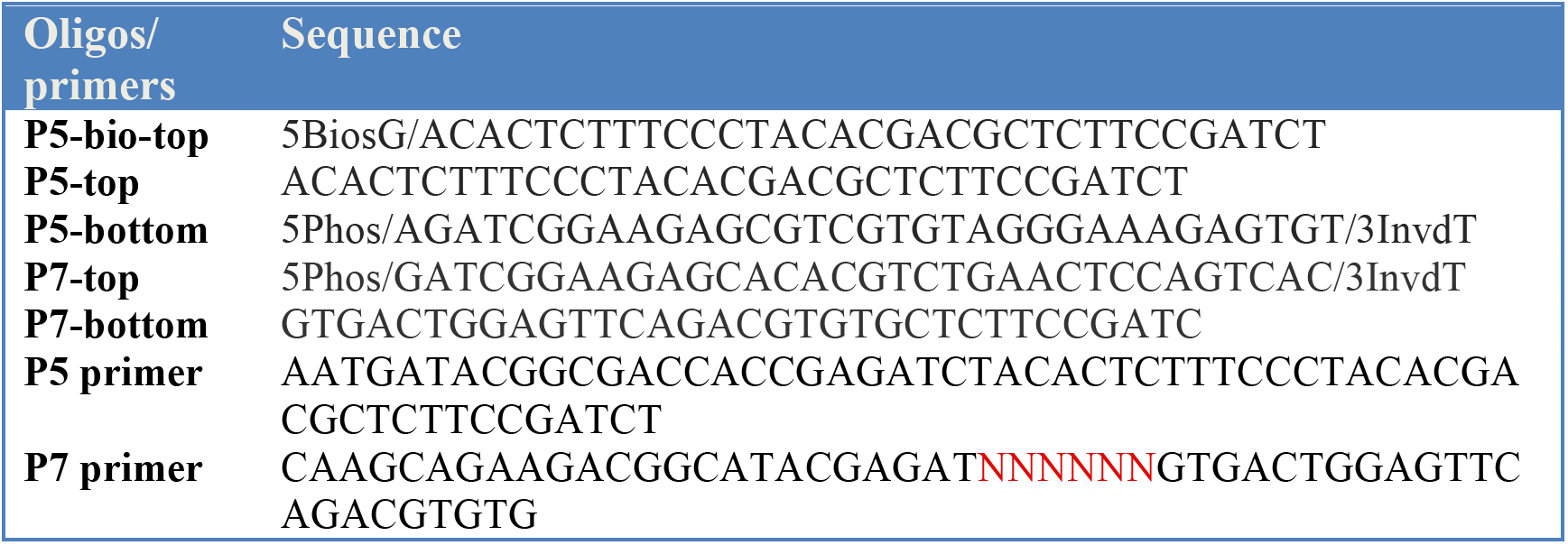
Oligonucleotide sequences used to prepare the adaptors and primers used to amplify the fragment library. NNNNNN denotes TruSeq barcode sequence

#### 5.1.2 Procedure

1. For each sample or time point to be analysed, prepare ten 2-mL conical tubes with 500 μL 1× TE.
2. Place one plug in each tube to rinse away the plug storage buffer.
3. Aspirate 1× TE and replace with 500 μL 1× S1 buffer. Equilibrate for 30 min. See note 3.
4. Repeat step 3 three times, so total equilibration time in 1× S1 buffer is 2 h.
5. Aspirate and replace with 500 ul 1× S1 buffer containing 9 U of S1 nuclease.
6. Place on ice for 15 min to allow the enzyme diffuse into the plugs.
7. Incubate at 37°C for 20 min.
8. Inactivate S1 nuclease by addition of EDTA pH 8.0 to a final concentration of 10 mM and incubate on ice for 15 min.
9. Rinse plugs with 500 μL 1× TE.
10. Equilibrate in 500 μL T4 polymerase reaction buffer, four times for 30 min each.
11. At this point prepare the P5 adaptor (biotinylated and optionally non-biotinylated), by mixing oligos P5-top (biotinylated and non-biotinylated, 100 μM) and P5-bottom (100 μM) at equimolar concentration, boiling for 5 min, and cooling at room temperature for at least 1 h.
12. Aspirate and add 500 μL T4 polymerase reaction buffer containing 30 U of T4 DNA polymerase.
13. Incubate on ice for 15 min, followed by 30 min incubation at 12°C.
14. Inactivate T4 polymerase by addition of EDTA pH 8.0 to a final concentration of 10 mM and incubate on ice for 15 min.
15. Aspirate buffer and rinse plugs with 1× TE.
16. Perform a quick spin and remove all buffer.
17. Incubate at 75°C for 20 min to fully inactivate T4 polymerase.
18. Allow plugs to gradually cool and solidify at room temperature.
19. Equilibrate in 250 μL 1× T4 ligase buffer, four times for 15 min each.
20. Aspirate 200 μL of the buffer leaving each plug immersed in ~50 μL 1× T4 DNA ligase buffer.
21. Add 1 μL of 50 μM adaptor and 1 μL of 2000 U/μL T4 DNA ligase to each plug.
22. Incubate at 16°C for ≥18 h.

### 5.2 Gel extraction and size selection

This step aims at releasing total DNA from the agarose plugs and further separating large genomic DNA fragments from residual unligated P5 adaptor. We found that including the size chromatography step reduced the background at later steps in this protocol (PCR amplification) so we deem it important (see note 4).

#### 5.2.1 Material, Solutions, and Reagents

1. GELase Agarose Gel-Digesting Preparation (Epicentre, currently discontinued, can be replaced with NEB β-agarase)
2. Phenol:Chloroform pH 8 (Amresco)
3. 3 M sodium acetate pH 5.5
4. 100 % ethanol
5. 70 % ethanol
6. 6× loading dye (NEB)
7. 10 mM Tris-HCl pH 7.5
8. 50× Tris-Acetate-EDTA (Fisher)
9. LMP agarose (Lonza SeaPlaque)
10. Scalpels or razor blades
11. Equipment for UV visualization and photography of DNA gel
12. 5 mg/mL ethidium bromide
13. Chroma spin +TE ‐1000 columns (Clontech)
14. Low-retention filter tips
15. Low-retention tubes: Siliconized G-tube microcentrifuge tubes, 1.5 mL (VWR)

#### 5.2.2 Procedure

1. Add 3 μL of 50× GELase buffer to each plug and incubate at 70°C until plug has thoroughly melted, approximately 5-10 min.
2. Pool plugs of the same sample together and equilibrate at 45°C for ≥ 5 min.
3. Add 1 unit GELase per plug (i.e., 10 U total) and incubate at 45°C for 30–45 min.
4. Add 1:1 ratio of phenol, mix by inversion and spin at max speed for 10 min.
5. Carefully transfer the aqueous phase to new tubes.
6. Add 0.1 volume of sodium acetate pH 5.5 and 2.5 volumes 100% ethanol, mix by inversion 5–6 times, and place at ‐20°C for ≥1 hr.
7. Centrifuge at full speed for 15 min and wash pellet once with 70% ethanol.
8. Dry pellet at room temperature for 15 min.
9. Solubilize in 120 μL 10 mM Tris-HCl pH 7.5.
10. Add 24 μL 6× loading dye and mix well by gently flicking the tube (see note 5).
11. Meanwhile prepare a 1% LMP 1× TAE agarose gel, containing 0.5 μg/mL ethidium bromide.
12. Carefully load the genomic DNA into the wells (see note 5) and subject to electrophoresis at 80 V/cm for 2 h.
13. Visualize the gel; a typical run is shown in Figure 5A. Excise the high molecular weight band migrating beneath the wells.
14. Weigh the gel slice, add GELase buffer to 1× final concentration and melt the gel slice thoroughly at 70°C.
15. After equilibration at 45°C for 5 min add 1 unit GELase for every 200 mg gel slice and incubate at 45°C for 30 min.
16. Add phenol in 1:1 ratio, mix by inversion, and spin at max speed for 10 min. Remove aqueous phase and place in fresh tubes.
17. Add 0.1 volumes of sodium acetate pH 5.5 and 2.5 volumes 100% ethanol, mix by inversion 5–6 times, and place at ‐20°C for ≥1 hr.
18. Centrifuge at full speed for 15 min and wash pellet once with 70 % ethanol.
19. Dry pellet at room temperature for 15 min.
20. Solubilize pellet in 100 μL 1× TE.
21. Pass each sample through Chroma Spin-1000 columns.
22. Repeat step 21 two more times.
23. Subject the final eluates to shearing according to the Covaris instrument protocol to DNA fragment sizes ranging between 200–500 bp. To verify the size range of the library, a dilution of the library is analyzed on a Bioanalyzer Instrument (Agilent Technologies) according to manufacturer’s instruction. An example is shown in Figure 5B.

**Figure 5.**
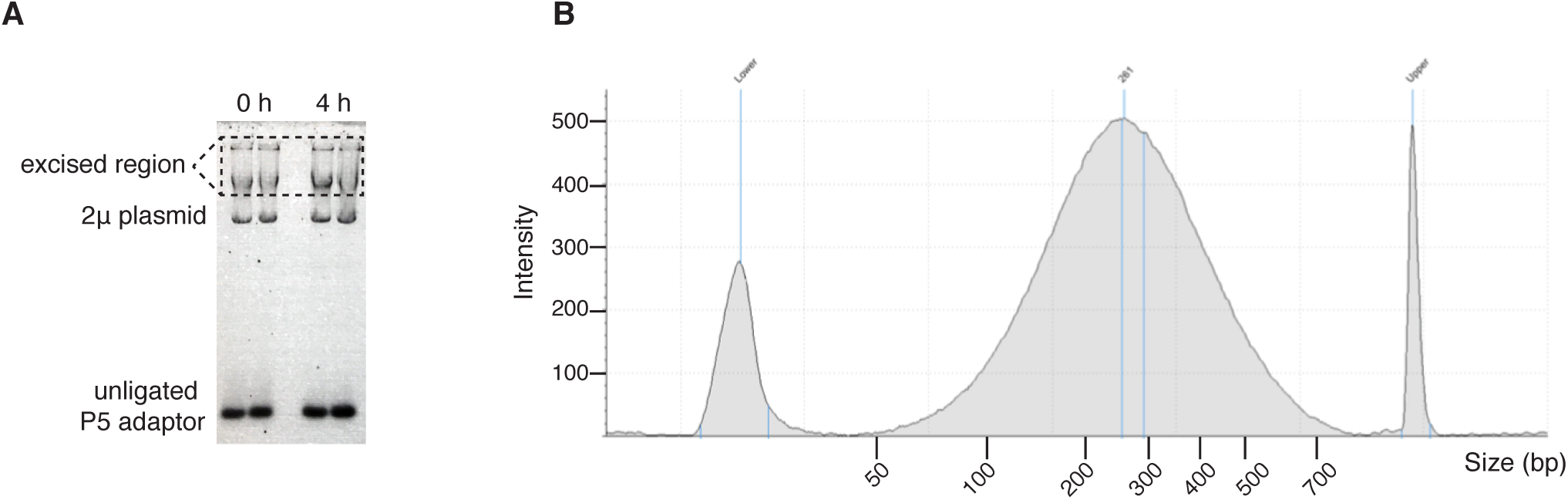
Gel electrophoresis to separate high molecular weight DNA from the unligated adaptor: A. Example of the agarose gel after electrophoresis. The boxed region is excised and treated with agarase to retrieve the DNA. B. Bioanalyzer (Agilent) lane profile showing the distribution of the genomic DNA after Covaris shearing (~100-700 bp).

### 5.3 Streptavidin purification, end repair, and second-end ligation

This step enriches for fragments containing the biotinylated P5 adaptor followed by ligation of the P7 adaptor. Upon capture, all subsequent steps up to PCR amplification can be performed on beads.

#### 5.3.1 Material, Solutions, and Reagents

1. Dynabeads M-280 Streptavidin, (ThermoFisher)
2. Magnet for capturing Dynabeads
3. 1× TE: 10 mM Tris, 1 mM EDTA
4. 2× B&W buffer: 10 mM Tris-HCl pH 7.5, 1 mM EDTA, 2 M NaCl
5. End-It DNA End-Repair Kit (Epicentre)
6. 1 M Tris-HCl pH 7.5
7. 10× T4 ligase reaction buffer (NEB)
8. T4 DNA ligase, 2000 U/μL (NEB)
9. P7-top oligo 100 μM, dissolved in 10 mM Tris-HCl pH 7.5, 50 mM NaCl, 1 mM
10. EDTA pH 8 (see table 1)
11. P7-bottom oligo 100 μM, dissolved in 10 mM Tris-HCl pH 7.5, 50 mM NaCl, 1 mM EDTA pH 8 (see table 1)
12. Low-retention filter tips
13. Low-retention tubes: Siliconized G-tube microcentrifuge tubes, 1.5 mL (VWR)

#### 5.3.2 Procedure

1. Vortex the beads to fully resuspend and pipette the appropriate volume in a fresh eppendorf tube (you will need 50 μL beads per sample).
2. Capture beads with magnet and wash twice with 1× TE and twice with 1× B&W (wash with at least double the volume of beads).
3. Capture beads and resuspend in 2× B&W, using double the volume withdrawn at step 1.
4. Aliquot 100 μL beads into fresh tubes and add equal volume (adjust with 1× TE if less than 100 μL) of sheared DNA sample.
5. Allow binding by incubating at 20°C for 30 min.
6. During the incubation prepare P7 adaptor: anneal P7 top and P7 bottom in 1:1 ratio, boil for 5 min and allow to cool down at room temperature for at least one hour.
7. Wash twice with 500 μL 1× B&W.
8. Wash twice with 500 μL 10 mM Tris-HCl pH 7.5.
9. During the last wash, prepare the end-repair reaction mix, using the End-it DNA Repair kit (Epicentre). Each reaction will be 100 μL, scale accordingly: 68 μL ddH_2_O, 10 μL 10× buffer, 10 μL dNTPs, 10 μL ATP, and 2 μL enzyme mix.
10. Resuspend beads in 100 μL reaction mix and incubate at 20°C for 45 min.
11. Capture beads and wash twice with 500 μL 1× B&W.
12. Wash twice with 500 μL 10 mM Tris-HCl pH 7.5.
13. Wash once with 100 μL 1× T4 ligase buffer.
14. Meanwhile prepare the ligation reaction mix; per reaction: 87.8 μL ddH_2_O, 10 μL 10× ligase buffer, 0.2 μL 2000U/ μL T4 ligase.
15. Resuspend beads in 98 μL of ligation reaction mix and add 2 μL 50 μM P7 adaptor.
16. Incubate at 20°C for 3-4 h.
17. Collect beads, wash three times with 500 μL 1× TE and three times with 500 μL 10 mM Tris-HCl.
18. Resuspend beads in 20 μL 10 mM Tris-HCl pH 7.5 and store at 4°C.

## 5.4 PCR amplification and size selection

### 5.4.1 Material, Solutions, and Reagents

1. PCR tubes (e.g., PCR 8-strips (Brandtech Scientific)).
2. Phusion High-Fidelity DNA Polymerase (Thermo Scientific)
3. 100 mM dNTPs set (Sigma Aldrich)
4. Barcoded sequencing primers for Illumina library (TruSeq Indexed Primer P7, TruSeq Universal Primer P5). See table 1.
5. Thermocycler
6. Equipment for UV visualizatio1n and photography of DNA gel
7. 7.5 M ammonium acetate
8. 5% Mini-Protean TBE gel (Biorad)
9. 1× TBE diluted from 10χ TBE (890 mM Tris, 890 mM boric acid, 20 mM EDTA)
10. 0.6 mL tubes (Fisher)
11. BD PrecisionGlide Single-use Needles 19 Gauge (Fisher)
12. Spin-X centrifuge Tube filter, 0.45 μm cellulose acetate (Corning costar)
13. Molecular weight markers (e.g., Low Molecular Weight Marker (NEB)).
14. Ethidium bromide
15. Platform shaker and tray for gel staining with ethidium bromide
16. Dedicated bottle of ethanol for ethanol precipitation
17. 10 mM Tris-HCl pH 8.0
18. Spin-X column: Corning Incorporated Costar Centrifugal Devices, cellulose acetate membrane (Fisher)
19. Scalpels or razor blades
20. Saran wrap
21. Thermomixer
22. Low-retention filter tips
23. Low-retention tubes: Siliconized G-tube microcentrifuge tubes, 1.5 mL (VWR)

#### 5.4.2 Procedure

1. Split each 20 μL bead sample into two PCR tubes, using 10 μL as template:

2. Run the PCR with the following conditions: 98°C for 30 sec; 16 cycles of 98°C for 10 sec, 65°C for 20 sec, and 72°C for 20 sec; and a final extension step at 72°C for 5 min.
3. Handle PCR products on a different bench from the one used to set up the reactions, and use designated pipettes and tips, as well as reagent aliquots reserved for PCR products. UV-irradiate both sides of the pipettes using 1200 J/m^2^ before use.
4. Combine the two reactions coming from the same sample into one 1.5 mL tube.
5. Capture beads and move the supernatant (100 μL) into fresh 1.5 mL tubes.
6. Precipitate the DNA by adding 50 μL 7.5 M ammonium acetate and 250 μL 100% ethanol.
7. Incubate at ‐20°C overnight.
8. Centrifuge at maximum speed for ≥30 min at 4°C.
9. Wash the pellet with cold 70% ethanol. A pellet should be clearly visible at this step. Centrifuge at maximum speed for 10 min at 4°C. Remove all supernatant, and perform a quick spin to gather residual ethanol and remove with a pipette. Air dry for 10-15 min.
10. Dissolve the pellets by adding 20 μL of 10 mM Tris-HCl pH 7.5.
11. Set up the 5% Mini-Protean TBE gel in the Biorad tank and add 1× TBE as running buffer.
12. Add 4 μL 6× DNA loading dye to each PCR product and load everything into one lane.
13. Run the gel at 200 V until just before the dye front migrates out of the gel (~30 min).
14. Stain the gel with 0.5 μg/mL ethidium bromide for 15 min and image the gel. Make sure to wipe all surfaces, and place the gel on Saran wrap (not directly on any surface) to avoid contamination. A smear spanning from 200-800 bp should be visible, with primer dimers migrating at ~100 bp. An example is shown in Figure 6A.
15. Set up a centrifuge device by punching a hole at the bottom of a 0.6 mL tube using a 19G1/2 needle, and placing it into an open 1.5 mL tube.
16. Using a UV transilluminator and wearing appropriate eye protection, cut out the 200-600 bp smeared region and transfer to the centrifuge device.
17. Centrifuge at maximum speed for 2 min. Discard the 0.6 mL tube.
18. Add 300 μL 10 mM Tris-HCl pH 7.5.
19. Elute overnight in a 37°C thermomixer with shaking at 800 rpm.
20. Transfer the gel-eluate mixture from one tube to the Spin-X column. Centrifuge at maximum speed for 2 min.
21. Add 150 μL 7.5 M ammonium acetate and 750 μL 100% ethanol to the ~300 μL eluate. Mix by inverting the tube. Precipitate at ‐20°C overnight.
22. Centrifuge at maximum speed for ≥30 min at 4°C.
23. Wash with 300 μL cold 70% ethanol. Centrifuge at maximum speed for 10 min at 4°C. Remove supernatant, then quickly spin and pipette out all remaining ethanol. Air dry for 10-15 min.
24. Dissolve the pellet in 20 μL 10 mM Tris-HCl pH 7.5.
25. To verify the size range of the library, a dilution of the library is analyzed on a Bioanalyzer Instrument (Agilent Technologies) according to manufacturer’s instruction. An example is shown in Figure 6B.
26. Proceed with next-generation sequencing (50 bp single-end reads), or store at ‐20°C until ready (see note 6).

| Component | Per reaction (μL) | Final Concentration |
| --- | --- | --- |
| Beads in 10 mM Tris-HCl | 10 |  |
| 5× Phusion HF buffer | 10 | 1 × |
| 10 mM dNTPs | 1 | 0.2 mM |
| Phusion High-Fidelity 2 U/ μL | 0.5 | 1 U |
| 10 μM primer P5 | 2 | 0.4 μM |
| 10 μM primer P7 (indexed) | 2 | 0.4 μM |
| ddH<sub>2</sub>O | 24.5 |  |
| Total volume | 50 |  |

**Figure 6.**
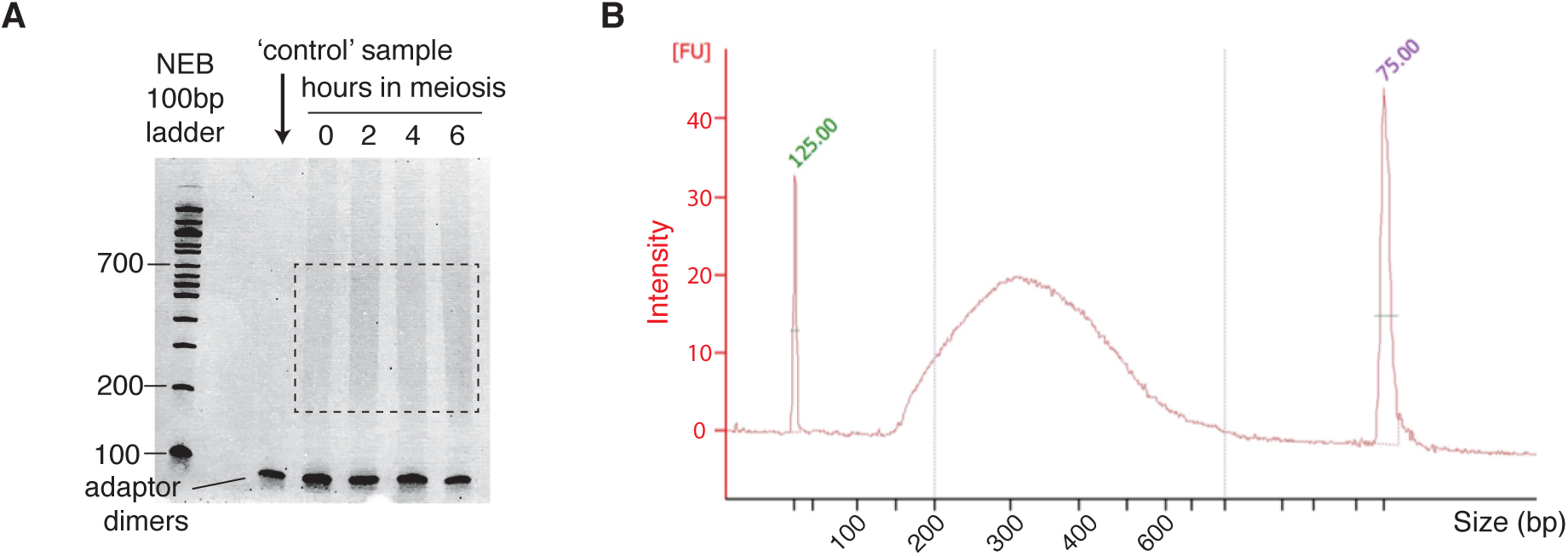
PCR (sequencing libraries) of a wild-type time course. A. Samples from different time points (0, 2, 4 and 6 h) were resolved on a 5% non-denaturing polyacrylamide gel. The first lane is a controlwhere the non-biotinylated P5 adaptor was used. The smear at 200-700 bp (brackets) indicates PCR-amplified adaptor-ligated fragments and is excised from the gel. The DNA is extracted and ethanol precipitated prior to next-generation sequencing. **B.** Bioanalyzer (Agilent) lane profile showing the S1-seq library size distribution (~150–600 bp).

## 6. Bioinformatics analysis

### 6.1 Mapping of reads

Mapping of the reads onto the S288c reference genome (SacCer2) is performed using the SHRiMP mapper (Rumble et al., 2009) (gmapper-ls) with arguments:

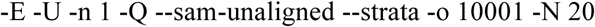

Before mapping, adaptor sequences are removed using fastx_clipper (http://hannonlab.cshl.edu/fastx_toolkit/) and a custom script. Code used for read processing and mapping is available online at https://github.com/soccin/S1-seq. After mapping, the reads are separated into unique and multiple-mapping sets, but only uniquely mapping reads are analyzed.

### 6.2 Curation of data

Before any analysis, maps are curated by masking 500 bp at the ends of each chromosome, because telomeres represent naturally occurring resected DNA ends and therefore give a high frequency of reads. We further mask regions that give a high frequency of meiotic DSB-independent reads (Mimitou et al., 2017). A table of mask coordinates is provided at https://github.com/soccin/S1-seq. Moreover, reads mapping to mitochondrial DNA or the 2μ plasmid are excluded. Each map is normalized to reads per million remaining mapped reads and then biological replicates are averaged.

### 6.3 Data analysis

Analyses can be performed using the R package. Our analyses use a hotspot list compiled from a combination of multiple independent wild-type Spo11-oligo maps (Mohibullah & Keeney, 2016). For quantitative and modeling analyses we use a subset of hotspots for which no other hotspot is located within 3 kb, and for which hotspot width is less than 400 bp (n=405). S1-seq reads of polarity opposite to expectation for resection endpoints are subtracted to correct for RI signal and any negative values that arise are set to NA.

## 7. Notes

1. To monitor the progression of cell divisions, collect a small volume (≥ 25 μL) of the sporulation culture at least every two hours (e.g., t = 0, 2, 4, 6, 8 h, and one at a later time point e.g., t = 24 h for *S. cerevisiae* SK1). Harvest cells and fix in 100 μL 40% ethanol in 0.1 M sorbitol. Add 1 μL 10 mg/mL DAPI (4′, 6-diamidino-2-phenylindole) and score mono-, bi‐ and tetranucleate cells by fluorescence microscopy. Fixed cells can be stored at 4°C until ready to count.
2. The cell suspension is very dense at this point, so avoid moving the cells from tube to tube. Instead, add solution 1 and LMP to the cells, either as a premix or sequentially and quickly mix and cast into the plugs.
3. Perform all buffer exchanges carefully — so as not to disrupt the plugs — by guiding the pipet tip to the side of the tube. Incubations can be performed with gentle shaking, on a horizontal platform mixer for example, but we do not find this necessary.
4. We noticed substantial DNA loss after the Chroma-Spin column step, but omitting it altogether resulted in the presence of background bands after PCR amplification. More recent S1-seq libraries in our laboratory replaced this step with sonicator shearing (200-500 bp), followed by SPRI size selection (e.g., Ampure XP beads). In this case the Covaris shearing step is also omitted.
5. The DNA solution at this point is extremely viscous because it contains high molecular weight fragments. Be patient: Pipet gently and slowly and mix by flicking the tube. Moreover, caution should be exerted while loading the samples onto the gel wells. Deposit sample inside the well slowly, avoiding air bubbles, and make sure the tip is no longer connected to the viscous solution when you remove it from the well.
6. The latest S1-seq libraries in our laboratory were submitted for paired-end sequencing and the improved mapping helped remove some of the non-specific signal, reducing the number of positions in the genome that need to masked.

## Acknowledgements

This work was supported by National Institutes of Health (NIH) grants R01 GM058673 and R35 GM118092 (to Sk) and a fellowship from the Helen Hay Whitney Foundation (to EM). MSKCC core facilities were supported by NIH grant P30 CA008748. SK is a Howard Hughes Medical Institute Investigator. Nicholas D. Socci (MSKCC Bioinformatics Core Facility) developed the sequence mapping pipeline and the MSKCC Integrated Genomics Operation core facility performed Covaris shearing and NGS. We are grateful to Shintaro Yamada for assistance in the bioinformatics analysis.

